# *Ir56d*-dependent fatty acid responses in *Drosophila* uncovers taste discrimination between different classes of fatty acids

**DOI:** 10.1101/2020.05.27.119602

**Authors:** Elizabeth B. Brown, Kreesha D. Shah, Justin Palermo, Manali Dey, Anupama Dahanukar, Alex C. Keene

## Abstract

Chemosensory systems are critical for evaluating the caloric value and potential toxicity of food prior to ingestion. While animals can discriminate between 1000’s of odors, much less is known about the discriminative capabilities of taste systems. Fats and sugars represent calorically potent and innately attractive food sources that contribute to hedonic feeding. Despite the differences in nutritional value between fats and sugars, the ability of the taste system to discriminate between different rewarding tastants is thought to be limited. In *Drosophila*, sweet taste neurons expressing the Ionotropic Receptor 56d (*IR56d*) are required for reflexive behavioral responses to the medium-chain fatty acid, hexanoic acid. Further, we have found that flies can discriminate between a fatty acid and a sugar in aversive memory assays, establishing a foundation to investigate the capacity of the *Drosophila* gustatory system to differentiate between various appetitive tastants. Here, we tested whether flies can discriminate between different classes of fatty acids using an aversive memory assay. Our results indicate that flies are able to discriminate medium-chain fatty acids from both short- and long-chain fatty acids, but not from other medium-chain fatty acids. Characterization of hexanoic acid-sensitive *Ionotropic receptor 56d* (Ir56d) neurons reveals broad responsive to short-, medium-, and long-chain fatty acids, suggesting selectivity is unlikely to occur through activation of distinct sensory neuron populations. However, genetic deletion of *IR56d* selectively disrupts response to medium chain fatty acids, but not short and long chain fatty acids. These findings reveal Ir56d is selectively required for fatty acid taste, and discrimination of fatty acids occurs through differential receptor activation within shared populations of neurons. These findings uncover a capacity for the taste system to encode tastant identity within a taste category.

## Introduction

Animals detect food primarily through taste and olfactory systems. Across phyla, there is enormous complexity in olfactory receptors and downstream processing mechanisms that allow for detection and differentiation between odorants (Keller et al., 2017; Nara, Saraiva, Ye, & Buck, 2011; Parnas, Lin, Huetteroth, & Miesenböck, 2013). By contrast, taste coding is thought to be simpler, with most animals possessing fewer taste receptors and a diminished ability to differentiate between tastants (Freeman & Dahanukar, 2015; Scott, 2018; Yarmolinsky, Zuker, & Ryba, 2009). Most early studies in different species have focused on characterization of a limited number of taste modalities largely defined by human percepts (sweet, bitter, sour, umami, salt), though there is growing appreciation that additional taste pathways are likely to influence gustatory responses and feeding (Chaudhari & Roper, 2010; Scott, 2018). Between studies of *Drosophila* and mammals, cells or receptors that are involved in sensing water, carbonation, fat, electrophiles, polyamines, metal ions, and ribonucleotides have been identified, suggesting a previously underappreciated complexity in the coding of tastants (Cameron, Hiroi, Ngai, & Scott, 2010; Kang et al., 2010; Mishra, Thorne, Miyamoto, Jagge, & Amrein, 2018; Y. V. Zhang, Ni, & Montell, 2013). Elucidating the underlying mechanisms of tastant detection can provide fundamental insight into the molecular and cellular basis of tastant recognition and taste processing.

In flies and mammals, tastants are sensed by dedicated gustatory receptors that are expressed in gustatory receptor neurons (GRNs) or taste cells respectively. In both systems, distinct subsets of taste sensory cells are activated by compounds belonging to distinct taste modalities such as sweet or bitter, and convey information to discrete areas of higher order brain structures (Vosshall & Stocker, 2007; Yarmolinsky et al., 2009; Yifeng Zhang et al., 2003). Given the conserved logic of taste processing, flies provide a powerful system for studying sensory processing and principles of taste circuit function (Freeman & Dahanukar, 2015; Scott, 2018; Yarmolinsky et al., 2009). Further, a number of genes and biochemical pathways that regulate feeding behavior are conserved across phyla (Vosshall & Stocker, 2007; Yarmolinsky et al., 2009). Notably, the gustatory system of *Drosophila* is amenable to *in vivo* Ca^2+^ imaging and electrophysiology, both of which can be coupled with robust behavioral assays that measure reflexive taste responses and food consumption (Wisotsky, Medina, Freeman, & Dahanukar, 2011). Taste neurons are housed in gustatory sensory structures called sensilla, which are located in the distal segments of the legs (tarsi), in the external and internal mouth organs (proboscis and pharynx), and in the wings. Each sensillum contains dendrites of multiple gustatory receptor neurons, each of which can be distinguished from the others based on its responses to various categories of tastants. Two main classes of non-overlapping gustatory neurons that have been identified are sweet-sensing and bitter-sensing neurons. Sweet-sensing GRNs promote feeding, whereas bitter-sensing GRNs act to deter (Marella et al., 2006; Thorne, Chromey, Bray, & Amrein, 2004). Both sweet and bitter GRNs express subsets of 68 G-protein-coupled gustatory receptors (GRs) (Clyne, Warr, & Carlson, 2000; Scott et al., 2001). In addition, the *Drosophila* genome encodes 66 glutamate-like Ionotropic Receptors (IRs), a recently identified family of receptors implicated in taste, olfaction, and temperature sensation (Benton, Vannice, Gomez-Diaz, & Vosshall, 2009; Rytz, Croset, & Benton, 2013). GRNs predominantly project to the subesophageal zone (SEZ), the primary taste center, but the higher order circuitry downstream of the SEZ contributing to taste processing is poorly understood (Flood et al., 2013; Marella, Mann, & Scott, 2012; Pool et al., 2014; Wang, Singhvi, Kong, & Scott, 2004). Determining how tastants activate GRNs that convey information to the SEZ, and how these signals are transmitted to higher order brain centers, is central to understanding the neural basis for taste and feeding.

In *Drosophila*, GRNs in the labellum and tarsi detect hexanoic acid (Pavel Masek & Keene, 2013). Mutation of Ionotropic receptor 56d (*IR56d*) disrupts hexanoic acid taste, implicating *IR56d* as a fatty acid receptor, or as part of a complex involved in fatty acid taste (Ahn, Chen, & Amrein, 2017; Sánchez-Alcañiz et al., 2018). *IR56d* is co-expressed with Gr64f (Ahn et al., 2017; Tauber et al., 2017), which broadly labels sweet GRNs (Dahanukar, Lei, Kwon, & Carlson, 2007; Jiao, Moon, Wang, Ren, & Montell, 2008; Slone, Daniels, & Amrein, 2007). *IR56d*-expressing GRNs are responsive to both sugars and fatty acids, suggesting that these neurons may respond to diverse appetitive substances including multiple classes of fatty acids (Tauber et al., 2017). Notably, overlapping populations of sweet GRNs that are responsive to different appetitive modalities and confer feeding behavior.

Is it possible that flies are capable of differentiating between tastants of the same modality, or is discrimination within a modality exclusively dependent on concentration? Taste discrimination can be assayed by training flies to pair a negative stimulus with a tastant, and determining whether the acquired aversion generalizes to another tastant (Keene & Masek, 2012; Pavel Masek & Scott, 2010). A previous study employing such experiments found that flies are unable to discriminate between different sugars (Pavel Masek & Scott, 2010). Conversely, we reported that flies can discriminate between sucrose (sugar) and hexanoic acid (fatty acid), revealing an ability to discriminate between appetitive stimuli of different modalities (Tauber et al., 2017). Here, we find that flies are capable of discriminating between different classes of fatty acids, despite broad tuning of fatty-acid sensitive neurons to short, medium and long chain FAs.

## Results

Sugars and medium chain fatty acids are sensed by an overlapping population of gustatory neurons, and flies can discriminate between these attractive tastants (Ahn et al., 2017; Tauber et al., 2017). To test whether flies are capable of discriminating within a single modality, we measured the ability of flies to discriminate between different types of fatty acids. We have used an appetitive taste memory assay in which an appetitive tastant is paired with bitter quinine, resulting in an associative memory that inhibits responses to the appetitive tastant (Pavel Masek, Worden, Aso, Rubin, & Keene, 2015). A modified version of this assay, in which training with one tastant is followed by testing with another, allows us to determine whether flies can discriminate between these tastants (Fig 1A). We first sought to determine whether flies are capable of differentiating between short (3C-5C), medium (6C-8C) and long (>9C) FAs. We found that flies that were trained with pairing of quinine and hexanoic acid (6C) exhibited PER to subsequent application of 5C fatty acids (Fig 1B). Thus, aversive memory to 5C was not formed by training with 6C, suggesting that flies can discriminate between these short- and medium-chain fatty acids. Similarly, flies trained with 6C did not generalize aversive memory to 9C, consistent with the idea that flies can also discriminate between medium- and long-chain fatty acids (Fig 1C). To rule out the possibility that flies are unable to form aversive taste memories to short- and long-chain fatty acids, we trained with 5C and found robust aversive taste memory, which did not generalize to 9C (Fig 1D). Together, these results suggest that flies are capable of distinguishing between short, medium and long-chain classes of fatty acids.

**Figure 1.**
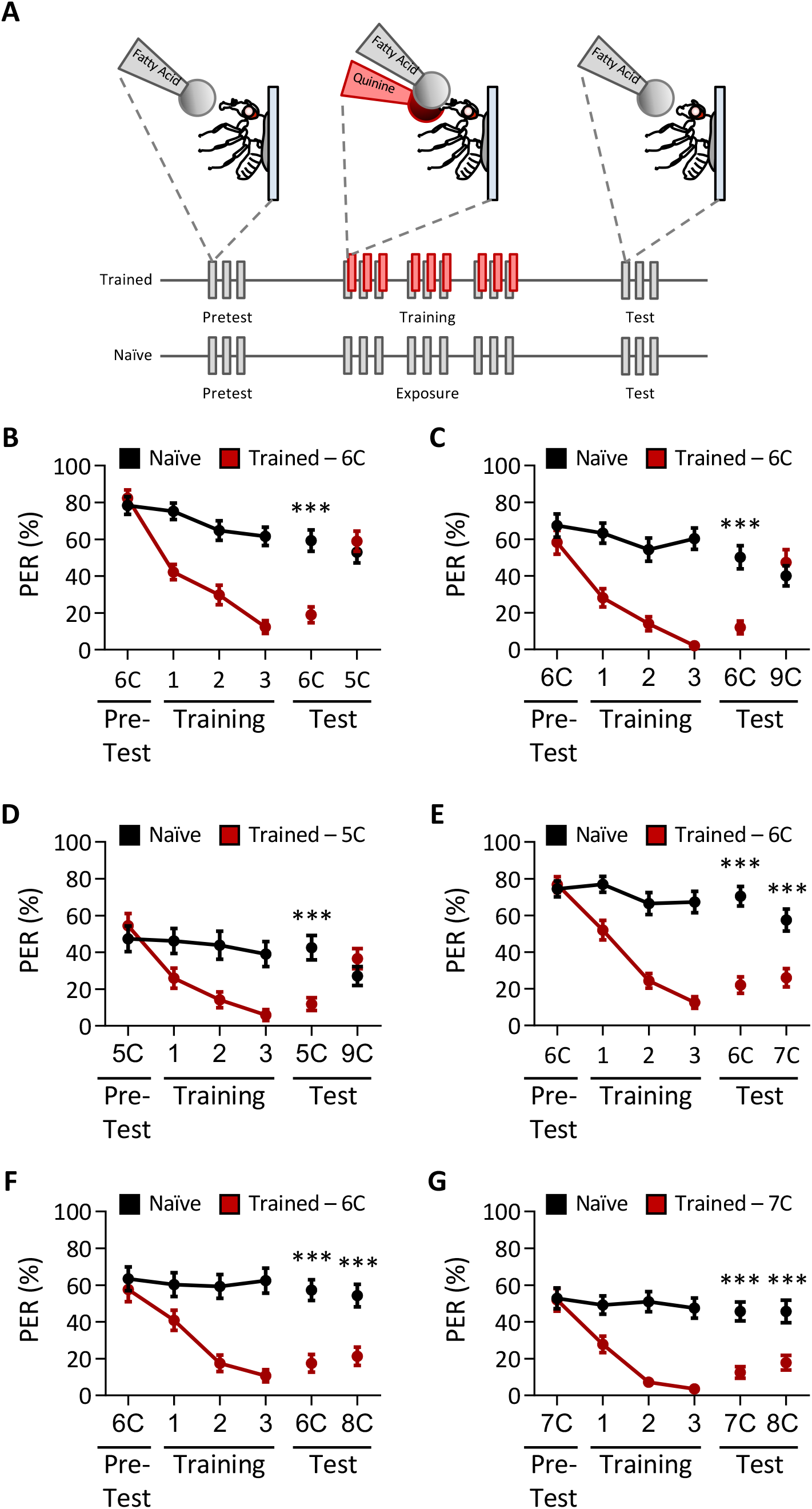
*Drosophila* can discriminate between short-, medium-, and long-chain fatty acids, but not among medium-chain fatty acids. **A** An aversive taste memory assay was used to assess FA taste discrimination. First, initial responses to a short-, medium-, or long-chain FA was assessed (Pretest). Next, flies were trained by pairing this FA with quinine (Training). PER in response to either the same or different FA was then tested in the absence of quinine (Test). In control experiments (Naïve), the same procedure was followed, but quinine was not applied to the proboscis. **B** The pairing of medium-chain hexanoic acid (6C) and quinine (red) results in a significant reduction in PER compared to naïve flies. After training, PER response to 6C was significantly lower in trained flies compared to naïve flies (*P*<0.0001), but there was no difference in PER to short-chain valeric acid (5C; *P*=0.6864). REML: F_1,80_ = 7.329, *P*=0.0003, with Sidak’s Test for multiple comparisons; N=40-42. **C** The pairing of medium-chain hexanoic acid (6C) and quinine (red) results in a significant reduction in PER compared to naïve flies. After training, PER response to 6C was significantly lower in trained flies compared to naïve flies (*P*<0.0001), but there was no difference in PER to long-chain nonanoic acid (9C; *P*=0.3346). REML: F_1,64_ = 6.296, *P*=0.0146, with Sidak’s Test for multiple comparisons; N=33. **D** The pairing of short-chain valeric acid (5C) and quinine (red) results in a significant reduction in PER compared to naïve flies. After training, PER response to 5C was significantly lower in trained flies compared to naïve flies (*P*=0.0014), but there was no difference in PER to long-chain nonanoic acid (9C; *P*=0.0789). REML: F_1,46_ = 2.721, *P*=0.0105, with Sidak’s Test for multiple comparisons; N=24. **E** The pairing of medium-chain hexanoic acid (6C) and quinine (red) results in a significant reduction in PER compared to naïve flies. After training, PER to both 6C and medium-chain heptanoic acid (7C) was significantly lower in trained flies compared to naïve flies (6C: *P*<0.0001; 7C: *P*<0.0001). REML: F_1,81_ = 45.88, *P*<0.0001, with Sidak’s Test for multiple comparisons; N=41-42. **F** The pairing of medium-chain hexanoic acid (6C) and quinine (red) results in a significant reduction in PER compared to naïve flies. After training, PER to both 6C and medium-chain octanoic acid (8C) was significantly lower in trained flies compared to naïve flies (6C: *P*<0.0001; 8C: *P*<0.0001). REML: F_1,65_ = 32.76, *P*<0.0001, with Sidak’s Test for multiple comparisons; N=33-34. **G** The pairing of medium-chain heptanoic acid (7C) and quinine (red) results in a significant reduction in PER compared to naïve flies. After training, PER to both 7C and medium-chain octanoic acid (8C) was significantly lower in trained flies compared to naïve flies (7C: *P*<0.0001; 8C: *P*<0.0001). REML: F_1,72_ = 33.67, *P*<0.0001, with Sidak’s Test for multiple comparisons; N=37.

To determine whether flies can discriminate between compounds within a single class of fatty acid, we tested the ability of flies to differentiate between different medium-chain fatty acids. We trained flies to associate 6C with quinine, while the medium-chain fatty acids 7C or 8C were not reinforced. In both cases, flies formed aversive memories to 6C, and this generalized to 7C and 8C, suggesting that flies cannot discriminate between different medium chain fatty acids (Fig 1E,F). To fortify these findings, we trained flies to 7C and measured the response to 8C. Again, flies formed aversive memory to 7C that was generalized to 8C (Fig 1G). Our findings reveal that flies cannot discriminate between different medium chain fatty acids, although they are able to discriminate medium-chain fatty acids from short- or long-chain fatty acids.

The short, medium, and long-chain fatty acids that we tested have distinctly different smells (Hallem & Carlson, 2006), raising the possibility that flies can discriminate between these compounds using a combination of olfactory and gustatory information. To exclude the effects of olfactory input, we surgically ablated the antennae, the maxillary palps, or both structures, and measured the ability of flies to discriminate between a representative tastant from each short-, medium-, or long-chain fatty acid (Fig 2A). All test groups were able to distinguish between sucrose and hexanoic acid, confirming that the ablation itself does not generally impact taste or memory formation (Fig 2B). Further, flies trained to a medium-chain fatty acid (6C) did not generalize aversion to short- (5C) or long-chain (9C) fatty acids, regardless of the absence of one or both olfactory organs (Fig 2C,D). Taken together, our findings reveal an ability of the taste system to encode the identity of different classes of fatty acids.

**Figure 2.**
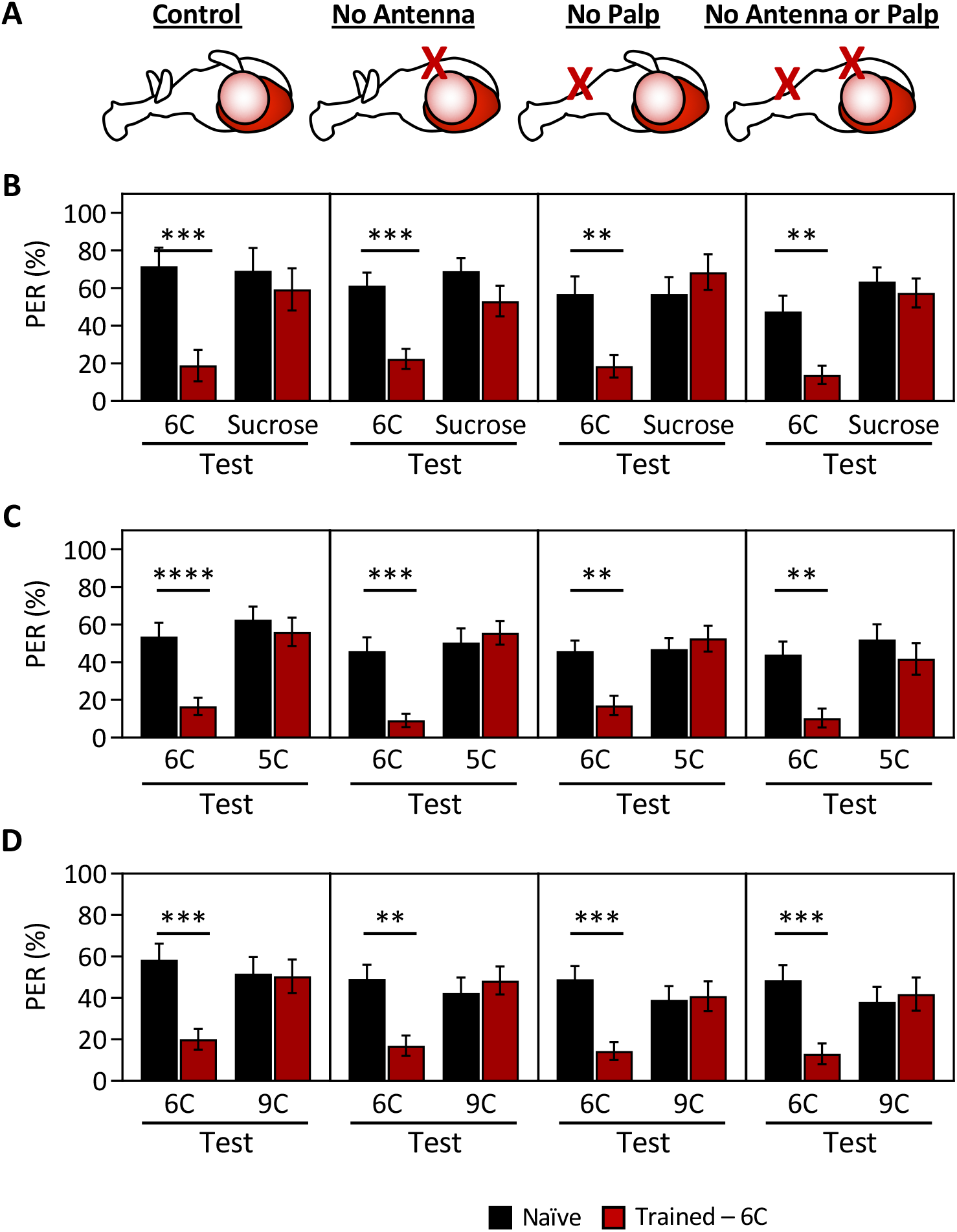
Ablation of chemosensory organs has no effect on the ability of *Drosophila* to discriminate between short-, medium-, and long-chain fatty acids. Aversive taste memory was measured as described in Figure 4A. Flies were trained by pairing medium-chain hexanoic acid (6C) with quinine (Training; See Figure S3) and then PER in response to either sucrose, shortchain valeric acid (5C), or long-chain nonanoic acid (9C) was measured in the absence of quinine (Test). **A** Aversive taste memory was measured in unmanipulated control flies (first panel), in flies without antennae (second panel), maxillary palps (third panel), or both antennae and maxillary palps (fourth panel). **B** For all ablation treatments, taste memory to medium-chain hexanoic acid (6C) was significantly lower in trained flies compared to naïve flies, but there was no difference in PER to sucrose. REML: F_1,86_ = 42.41, *P*<0.0001, with Sidak’s Test for multiple comparisons; N=13-26. **C** For all ablation treatments, taste memory to 6C was significantly lower in trained flies compared to naïve flies, but there was no difference in PER to short-chain valeric acid (5C). REML: F_1,103_ = 51.87, *P*<0.0001, with Sidak’s Test for multiple comparisons; N=19-31. **D** For all ablation treatments, taste memory to 6C was significantly lower in trained flies compared to naïve flies, but there was no difference in PER to long-chain nonanoic acid (9C). REML: F_1,97_ = 11.47, *P*=0.0010, with Sidak’s Test for multiple comparisons; N=22-27.

The finding that flies cannot discriminate between medium chain fatty acids raises the possibility that *IR56d* is required for the taste of medium-chain fatty acids, but not short and long-chain fatty acids. To determine whether *IR56d*-expressing GRNs mediate taste perception to other classes of fatty acids, we silenced *IR56d*-expressing neurons using the synaptobrevin cleavage peptide tetanus toxin light chain (TNT) (Sweeney, Broadie, Keane, Niemann, & Kane, 1995) and measured proboscis extension response (PER) to multiple classes of fatty acids, including short-, medium-, and long-chain fatty acids (Figure 3A). To control for any non-specific effects of TNT, we compared PER in flies with silenced *IR56d* GRNs (*IR56d*-GAL4>UAS-TNT) to flies expressing the inactive variant of TNT in *IR56d*-expressing GRNs (*IR56d*-GAL4>UAS-impTNT). Consistent with previous findings (Tauber et al., 2017), we observed no effect of silencing *IR56d*-expressing neurons on PER to sucrose (Figure 3B). Next, we measured PER to a panel of saturated FAs ranging from 4C (butanoic acid) to 10C (decanoic acid) in length (Figure 3C). Control flies exhibited a robust PER to all seven fatty acids, revealing that at least at a 1% concentration, many diverse classes of fatty acids can trigger this behavioral response. To determine whether *IR56d* is generally required for detection of fatty acids, or selectively required for sensing hexanoic acid, we next measured PER in flies with *IR56d*-expressing neurons silenced. Silencing *IR56d*-expressing neurons significantly reduced PER to the three medium chain fatty acids (6C, 7C, and 8C). Conversely, there was no difference in PER between control and *IR56d*-silenced flies in response to short chain (4C and 5C) and long-chain (9C and 10C) fatty acids. Therefore, *IR56d*-expressing neurons are required for medium-chain fatty acid taste perception, but are dispensable for responses to both short- and long-chain fatty acids.

**Figure 3.**
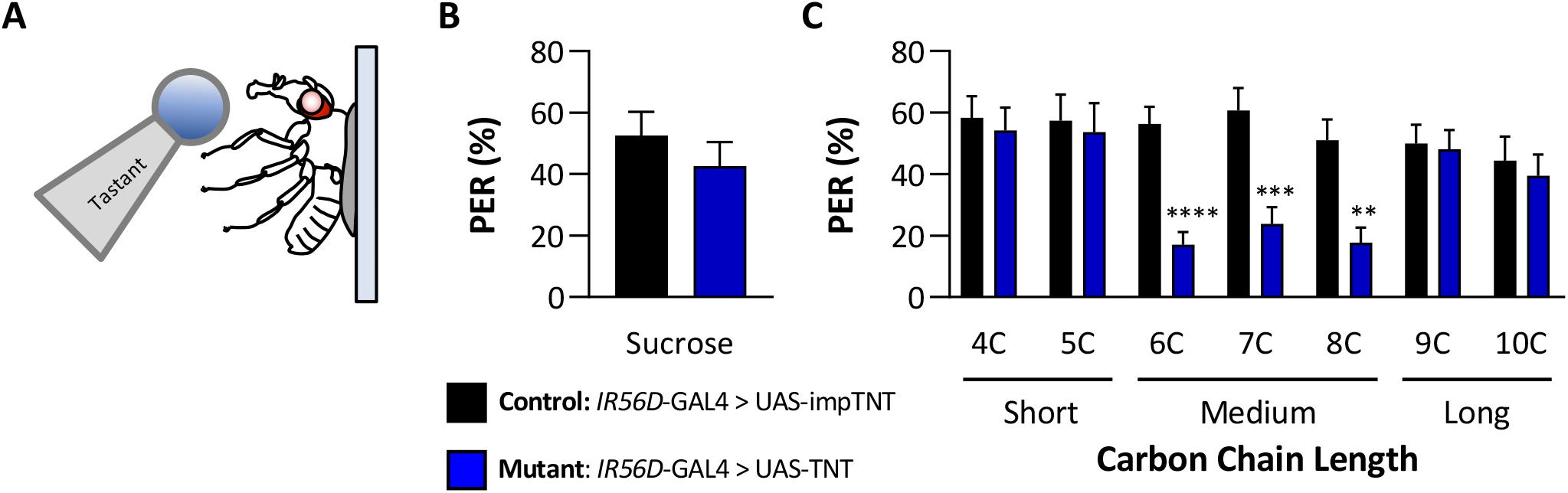
Silencing *IR56D*-expressing neurons reduces taste perception to medium chain fatty acids. **A** Proboscis extension response (PER). PER was measured in female flies after 48 hrs of starvation. Either sucrose or fatty acid was applied to the fly’s labellum for a maximum of two seconds and then removed to observe proboscis extension reflex. **B** Blocking synaptic release by genetic expression of light-chain tetanus toxin (UAS-TNT) in *IR56D*-expressing neurons has no effect on PER to sucrose compared to control flies expressing an inactive form of tetanus toxin (UAS-impTNT). Mann Whitney Test: U = 595, *P*=0.8410; N=35. **C** Silencing *IR56D*-expressing neurons significantly reduces PER to medium chain fatty acids (6C-8C), but has no effect on PER to either short- (4C,5C) or long-chain fatty acids (9C,10C). REML: F_1,406_ = 25.03, *P*<0.0001, with Sidak’s Test for multiple comparisons; N=24-45.

To directly assess whether the *IR56d* receptor mediates responses to medium chain fatty acids, we used the CRISPR/Cas9 system to generate an *IR56d* allele in which a GAL4 element is inserted into the *IR56d* locus (*IR56d*^GAL4^; Figure 4A), thereby allowing expression of UAS-transgenes under the control of the *IR56d* promoter. To confirm that the GAL4 knock-in element is indeed expressed in *IR56d* neurons, we generated flies carrying both UAS-mCD8:GFP and the *IR56d*^GAL4^ allele (*IR56d*^GAL4^>UAS-mCD8:GFP) and mapped the expression of GFP. Consistent with previous findings, we found GFP expression in labellar neurons that projected axons to both the taste peg and sweet taste regions of the SEZ (Figure 4B-E; (Koh et al., 2014; Tauber et al., 2017). In agreement with previous findings from genetic silencing of *IR56d*-expressing neurons, PER to sucrose did not differ between *IR56d*^GAL4^ and control flies (Figure 4F), suggesting that *IR56d*^GAL4^ is dispensable for response to sucrose. To examine the role of *IR56d* in fatty acid taste, we measured PER to fatty acids ranging from 4C to 10C in length (Figure 4G). Consistent with the results of *IR56d*-silenced flies, PER to medium chain fatty acids was disrupted in *IR56d*^GAL4^ flies (6C-8C), whereas PER to short- (4C and 5C) and long-chain fatty acids (9C and 10C) was not affected. Flies that were heterozygous for the *IR56d* deletion (*IR56d*^GAL4^/+) exhibited similar responses to those of control flies for all tastants measured. The observed decrease in PER to medium chain fatty acids was rescued by transgenic expression of *IR56d* in the *IR56d*^GAL4^ mutant background (*IR56d*^GAL4^; UAS-*IR56d*/+), confirming that the behavioral deficit of *IR56d*^GAL4^ flies is in fact due to loss of *IR56d* function. Therefore, *IR56d* appears to be selectively required for taste sensing of medium-chain fatty acids.

**Figure 4.**
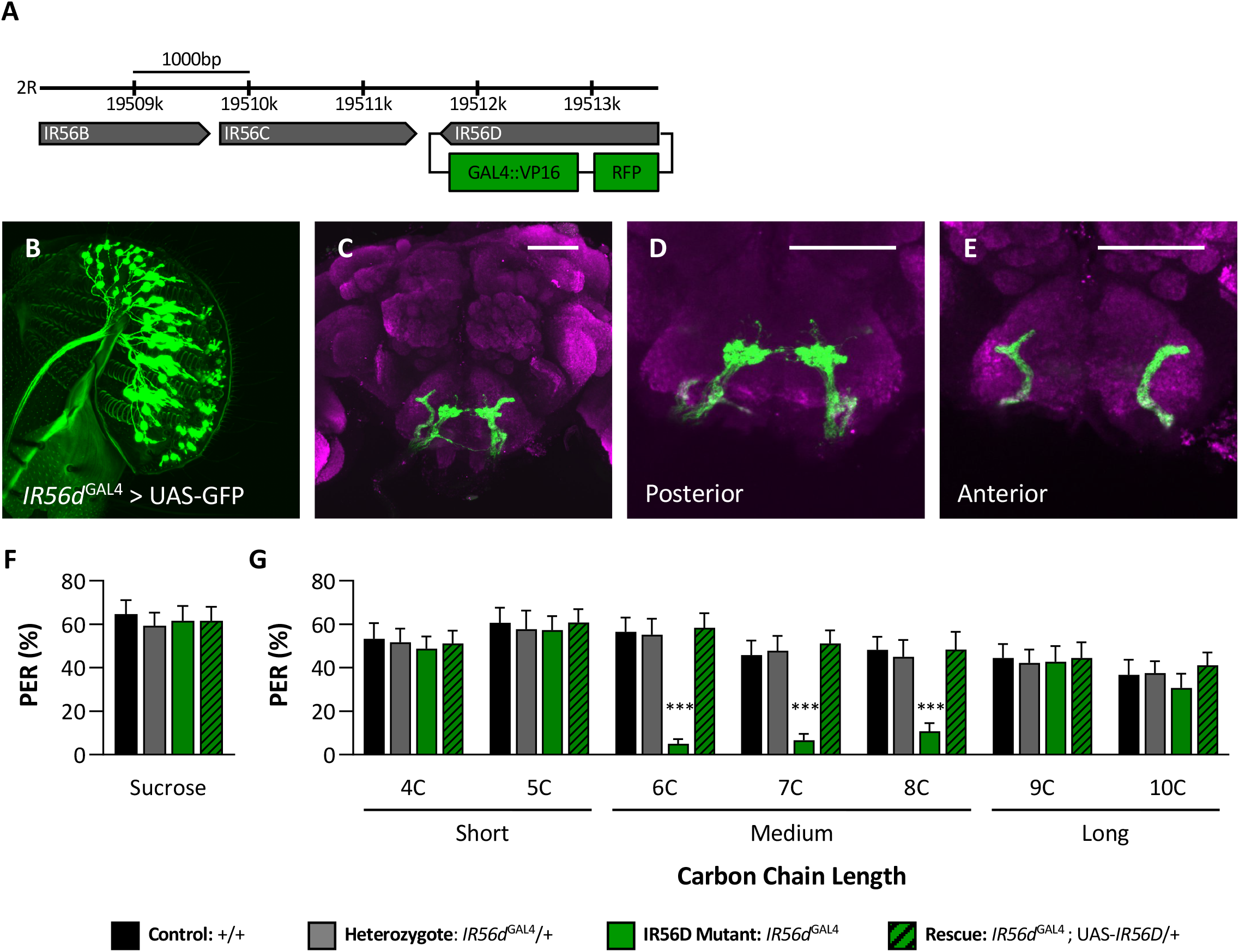
*IR56D* mediates taste perception to medium-chain fatty acids. **A** *IR56d*^GAL4^ was generated using the CRISPR/Cas9 system. In *IR56d*^GAL4^ flies, the *IR56D* gene was replaced by GAL4 and RFP elements (red boxes). The relative location and orientation of genes in the region are represented as gray arrows. **(B-E)** Expression pattern of *IR56d*^GAL4^ is visualized with GFP. *IR56D*-expressing neurons are located on the **(B)** labellum and project to the **(C)** subesophagael zone of the brain. Distinct regions of projection include the **(D)** posterior and **(E)** anterior subesophagael zones. Background staining is NC82 antibody (magenta). Scale bar = 50μm. **F** Sucrose taste perception is similar in control and *IR56d*^GAL4^ mutant flies. Kruskal-Wallis Test: H = 0.1758, *P*=0.9814, with Dunn’s Test for multiple comparisons; N=33-40. **G** The *IR56d*^GAL4^ flies have reduced PER to medium-chain fatty acids (6C-8C) relative to control, *IR56d*^GAL4^ heterozygotes, and *IR56d*^GAL4^ rescue flies. However, all genotypes respond similarly to both short- and long-chain fatty acids (4C,5C; 9C,10C). REML: F3,850 = 17.80, *P*<0.0001, with Tukey Test for multiple comparisons; N=28-40.

In previous work we found that *IR56d*-expressing neurons are activated by both sucrose and hexanoic acid (Tauber et al., 2017). To determine whether other classes of fatty acids can also activate these neurons, and if so, whether their activity is dependent on *IR56d*, we measured Ca^2+^ responses to a panel of tastants. We expressed the Ca^2+^ sensor GCaMP6.0 under the control of *IR56d*^GAL4^ and measured tastant-evoked activity (Figure 5A-D). In flies heterozygous for *IR56d*^GAL4^, the labeled neurons were activated by sucrose and all fatty acids tested, which ranged from 4C-10C (Figure 5E). Thus, *IR56d* neurons respond to diverse appetitive stimuli. Flies with a deletion of *IR56d*^GAL4^ (*IR56d*^GAL4^; *UAS*-GCaMP6.0) lacked responses exclusively to medium chain fatty acids (6C-8C), while responses to short- (4C and 5C) and long-chain fatty acids (9C and 10C) remained intact (Figure 5F). Consistent with the rescue of behavioral defects, inclusion of an *IR56d* rescue transgene (*IR56d*^GAL4^; *UAS-GCaMP6.0/UAS-IR56d*) restored the physiological response to medium chain fatty acids (Figure 5G). Quantification of the responses to all tastants confirmed that Ca^2+^ responses to 6C-8C fatty acids are disrupted in *IR56d*^GAL4^ flies, and restored to levels observed in control flies by expression of *IR56d* (Figure 5H). Overall, these results demonstrate that at both behavioral and physiological levels, *IR56d*^GAL4^ is required for taste responses to medium chain fatty acids. Therefore, these findings support the notion that medium chain fatty acids are detected through a shared sensory channel, allowing flies to distinguish medium chain from short or long-chain, but not between different medium-chain fatty acids.

**Figure 5.**
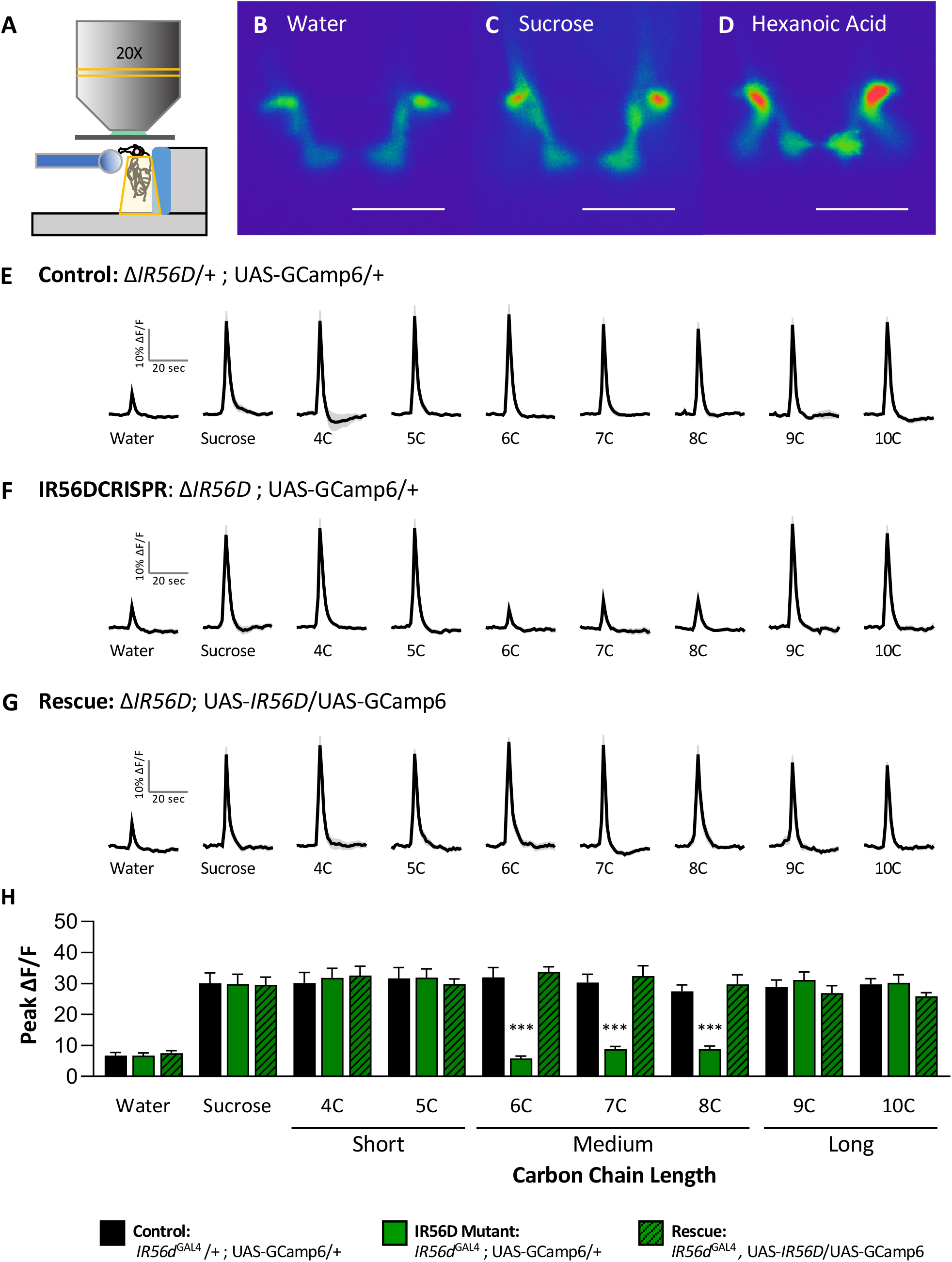
Neuronal activity of *IR56d*^GAL4^ mutant flies is reduced in response to medium-chain fatty acids. **A** Diagram of live-imaging experimental protocol. A tastant is applied to the proboscis while florescence is recorded simultaneously. **(B-D)** Representative pseudocolor images of calcium activity of the posterior projections of *IR56D* neurons in response to water **(B)**, 10mM sucrose **(C)**, or 1% hexanoic acid **(D)**. Scale bar = 50μm. **(E-G)** Activity traces of the posterior projections of *IR56D* neurons in response to each tastant in the **(E)** *IR56d*^GAL4^ heterozygote controls, **(F)** *IR56d*^GAL4^ mutants, and **(G)** *IR56d*^GAL4^ rescue flies. The shaded region of each trace indicates ±SEM. **H** Average peak change in fluorescence for data shown in E-G. Neuronal responses of to medium-chain fatty acids (6C-8C) are significantly reduced in *IR56d*^GAL4^ mutants compared to *IR56d*^GAL4^ heterozygote controls and *IR56d*^GAL4^ rescue flies. All genotypes respond similarly to both short- and long-chain fatty acids (4C,5C; 9C,10C), was well as to water and sucrose. Two-way ANOVA: F2,256 = 23.67, *P*<0.0001, with Sidak’s Test for multiple comparisons; N=8-14.

## Discussion

Receptors for sweet and bitter taste have been well defined in both flies and mammals (Carleton, Accolla, & Simon, 2010; Hallem, Dahanukar, & Carlson, 2006; Scott, 2018), but less is known about detection of fats. Previous studies identified *IR56d* as a receptor for hexanoic acid and carbonation (Ahn et al., 2017; Sánchez-Alcañiz et al., 2018). Our findings suggest that *IR56d* is selectively involved in responses to medium-chain fatty acids, including 6C, 7C, and 8C fatty acids, and dispensable for responses to shorter and longer-chain fatty acids. Such receptor specificity for different classes of fatty acids based on chain length has not been documented in other systems. In flies, both sugars and fatty acids activate neurons that coexpress the receptors *Gr64f* and *IR56d*. The finding that short- and long-chain fatty acids also activate *IR56d*-expressing neurons posits that additional fatty acid receptors are present in these neurons. Previously, we found that deletion of PLC signaling selectively impairs hexanoic acid response while leaving sweet taste intact, raising the possibility that activation of distinct intracellular signaling pathways could serve as a mechanism for discrimination of sucrose and hexanoic acid (Pavel Masek & Keene, 2013; Tauber et al., 2017). Examining whether or not short- and long-chain fatty acids also signal through phospholipase C may provide insight into whether signaling mechanisms are shared between different fatty acid receptors expressed in *IR56d* neurons.

Our aversive taste memory assay confirmed previous findings that flies can discriminate between sugars and fatty acids (Tauber et al., 2017), and led to the surprising observation that flies can distinguish between different classes of fatty acids. This contrasts with the results of a previous study that applied a similar assay and found that flies were unable to discriminate between different sugars or bitter compounds (Kirkhart & Scott, 2015). One possibility is that this is due to differences in fatty acid detection, which is dependent on IRs, and sweet and bitter tastant detection, which relies on GRs (Chen & Dahanukar, 2020). These results suggest the ability of the *Drosophila* the taste system to discriminate may be more like the olfactory system than previously appreciated. Flies are able to distinguish between many different odorants, likely due to the complexity of olfactory coding at the level of the receptor as well as in the antennal lobe (Amin & Lin, 2019; Cognigni, Felsenberg, & Waddell, 2018; Guven-Ozkan & Davis, 2014). However, flies can also discriminate between odorants sensed by a single olfactory receptor, suggesting that temporal coding also plays a role in discrimination (DasGupta & Waddell, 2008). It is possible that similar mechanisms underlie discrimination between different classes of fatty acid tastants.

The *Drosophila* genome encodes 66 Ionotropic Receptors (IRs), which comprise a recently identified family of receptors implicated in taste, olfaction, and temperature sensation (Benton et al., 2009; Rytz et al., 2013). Ionotropic receptors are involved in the detection of many different tastants, and function as heteromers that confer sensory specificity (Rytz et al., 2013; van Giesen & Garrity, 2017). While *IR56d* expression is restricted to a subset of sweet taste neurons, it likely functions in a complex with *IR25a* and *IR76b*, all three of which are required fatty acid taste (Ahn et al., 2017; Sánchez-Alcañiz et al., 2018). Other tastants whose responses are mediated by IR receptors are also likely to be detected by IR complexes. For example, roles for *IR25a, IR62a* and *IR76b* have been described for Ca^2+^ taste (Thakur, Kim, Poudel, Montell, & Lee, 2017). The broad degree of co-expression of IRs in the brain and periphery can provide candidates for those involved in detecting short- and long-chain fatty acids.

The identification of taste discrimination between different classes of fatty acids provides the opportunity to identify how different tastants are encoded in the brain, and how these circuits are modified with experience. Although projections of primary taste neurons to the SEZ have been mapped in some detail, little is known about connectivity with downstream neurons and whether sensory neurons activated by different appetitive tastants can activate different downstream circuits. Recent studies have identified a number of interneurons that modify feeding, including IN1, a cholinergic interneuron activated by sucrose (Yapici, Cohn, Schusterreiter, Ruta, & Vosshall, 2016), E564 neurons that inhibit feeding (Mann, Gordon, & Scott, 2013), and *Fdg* neurons that are required for sucrose-induced feeding (Flood et al., 2013). Future work can investigate whether these, and other downstream neurons, are shared for fatty acid taste. Previous studies have found that incoming sensory information is selectively modulated within the antennal lobe in accordance with feeding state (Chu, Chui, Mann, & Gordon, 2014; LeDue et al., 2016). It will be interesting to determine if similar modulation promotes differentiation of sugars and fatty acids, which are sensed by shared gustatory neurons. Large-scale brain imaging has now been applied in flies to measure responsiveness to different tastants (Harris, Kallman, Mullaney, & Scott, 2015), and a comparison of brain activity patterns elicited by different classes of fatty acids may provide insight into differences in their sensory input and processing.

All experiments in this study tested flies under starved conditions, which is necessary to elicit the PER that is used as a behavioral readout of taste acceptance. However, responses to many tastants and odorants are altered in accordance with feeding state (LeDue et al., 2016; Root et al., 2008). For example, the taste of acetic acid is aversive to fed flies but attractive to starved flies, revealing a hunger-dependent switch (Devineni, Sun, Zhukovskaya, & Axel, 2019). Similarly, hexanoic acid activates both sweet and bitter sensing taste neurons, and the activation of bitter taste neurons is dependent on different receptors from those involved in the appetitive response (Ahn et al., 2017). Further, hunger enhances activity in sweet taste circuits, and suppresses that of bitter taste circuits, providing a mechanism for complex state-dependent modulation of response to tastants that activate both appetitive and deterrent neurons (Inagaki, Panse, & Anderson, 2014; LeDue et al., 2016).

The neural circuits that are required for aversive taste memory have been well defined for sugar, yet little is known about how fatty acid taste is conditioned. The pairing of sugar with bitter quinine results in aversive memory to sugar. Optogenetic activation of sweet taste neurons, which are activated by both sugar and fatty acids, in combination with quinine presentation is sufficient to induce sugar avoidance, suggesting that aversive taste memory does not depend on post-ingestive feedback (Keene & Masek, 2012). Further studies have elucidated that aversive taste memories are dependent on mushroom body neurons that form the gamma and alpha lobes, the PPL1 cluster of dopamine neurons, and alpha lobe output neurons, revealing a circuit regulating taste memory that differs from that controlling appetitive olfactory memory (Kirkhart & Scott, 2015; P Masek, Worden, Aso, Rubin, & Keene, 2015). It will be interesting to determine whether shared components regulate conditioning to fatty acids, or whether distinct mushroom body circuits regulate sweet taste and fatty acid conditioning. Further, examination of the central brain circuits that regulate aversive taste conditioning to different classes of fatty acids will provide insight into how taste discrimination is processed within the brain.

## Materials and Methods

### *Drosophila* stocks and maintenance

Flies were grown and maintained on standard food media (Bloomington Recipe, Genesee Scientific, San Diego, CA). Flies were housed in incubators (Powers Scientific, Warminster, PA, USA) on a 12:12 LD cycle at 25°C with humidity of 55-65%. The following fly strains were ordered from the Bloomington Stock Center: *w*^1118^ (#5905; (Levis, Hazelrigg, & Rubin, 1985)); *IR56D*-GAL4 (#60708; (Koh et al., 2014)), UAS-impTNT (#28840; (Sweeney et al., 1995)), UAS-TNT (#28838; (Sweeney et al., 1995)), UAS-GFP (#32186; (Pfeiffer et al., 2010)); UAS-GCaMP5 (#42037; (Akerboom et al., 2012)). UAS-Ir56d was generated using *Ir56d* cDNA, amplified with primers that generated a NotI-KpnI fragment that was cloned in the pUAS vector. The *IR56d*^GAL4^ line was generated by WellGenetics (Taipei City, Taiwan) using the CRISPR/Cas9 system to induce homology-dependent repair. At the gRNA target site, a dsDNA donor plasmid was inserted containing a GAL4::VP16 and RFP cassette. This line was generated in the *w*^1118^ genetic background and was validated by PCR and sequencing. All lines were backcrossed to the *w*^1118^ fly strain. For all experiments, mated female flies aged 7-to-9 days were used. For ablation experiments, the antenna and/or maxillary palp were removed two days post-eclosion.

### Reagents

The following fatty acids were obtained from Sigma Aldrich (St Louis, MO, USA): butyric acid (4C; #B103500), valeric acid (5C; #240370), hexanoic acid (6C; #21530), heptanoic acid (7C; #75190), octanoic acid (8C; #O3907), nonanoic acid (9C; #N5502), and decanoic acid (10C; #C1875). All fatty acids were tested at a concentration of 1% and were dissolved in water. Quinine hydrochloride was also obtained from Sigma Aldrich (#Q1125), while sucrose was purchased from Fisher Scientific (#FS S5-500; Hampton, New Hampshire, USA).

### Immunohistochemistry

Brains were prepared as previously described (Kubrak, Lushchak, Zandawala, & Nässel, 2016). Briefly, brains of 7-9 day-old female flies were dissected in ice-cold PBS and fixed in 4% formaldehyde, PBS, and 0.5% Triton-X for 30 minutes at room temperature. Brains were rinsed 3X with PBS and 0.5% Triton-X (PBST) for 10 minutes at room temperature and then incubated overnight at 4°C. The next day, brains were incubated in primary antibody (1:20 mouse nc82; Iowa Hybridoma Bank; The Developmental Studies Hybridoma Bank, Iowa City, Iowa, USA) diluted in 0.5% PBST at 4°C for 48 hrs. Next, the brains were rinsed 3X in 0.5% PBST 3X 10 minutes at room temperature and placed in secondary antibody (1:400 donkey anti-mouse Alexa Fluor 647; #A-31571; ThermoFisher Scientific, Waltham, Massachusetts, USA) for 90 minutes at room temperature. The brains were again rinsed 3X in PBST for 10 min at room temperature and then mounted in Vectashield (VECTOR Laboratories, Burlingame, CA). Brains were imaged in 2μm sections on a Nikon A1R confocal microscope (Nikon, Tokyo, Japan) using a 20X oil immersion objective. Images presented as the Z-stack projection through the entire brain and processes using ImageJ2 (Tauber et al., 2017).

### Proboscis Extension Response

Female flies were starved for 48 h prior to each experiment and then PER was measured as previously described (Pavel Masek & Keene, 2013; Tauber et al., 2017). Briefly, flies were anesthetized on CO_2_ and then restrained inside of a cut 200 μL pipette tip (#02-404-423; Fisher Scientific) so that their head and proboscis were exposed while their body and tarsi remain restrained. After a 60 min acclimation period in a humidified box, flies were presented with water and allowed to drink freely until satiated. Flies that did not stop responding to water within 5 minutes were discarded. A wick made of Kimwipe (#06-666; Fisher Scientific) was placed partially inside a capillary tube (#1B120F-4; World Precision Instruments; Sarasota, FL) and then saturated with tastant. The saturated wick was then manually applied to the tip of the proboscis for 1-2s and proboscis extension reflex was monitored. Only full extensions were counted as a positive response. Each tastant was presented a total of three times, with 1 min between each presentation. PER was calculated as the percentage of proboscis extensions divided by the total number of tastant presentations. For example, a fly that extends its proboscis twice out of the three presentation will have a PER response of 66%.

### *In vivo* calcium imaging

Female flies were starved for 48 h prior to imaging, as described (Tauber et al., 2017). Flies were anaesthetized on ice and then then restrained inside of a cut 200 μL pipette tip so that their head and proboscis were accessible, while their body and tarsi remain restrained. The proboscis was manually extended and then a small amount of dental glue (#595953WW; Ivoclar Vivadent Inc.; Amherst, NY) was applied between the labium and the side of the pipette tip, ensuring the same position throughout the experiment. Next, both antennae were removed. A small hole was cut into a 1 cm^2^ piece of aluminum foil and then fixed to the fly using dental glue, creating a sealed window of cuticle exposed. Artificial hemolymph (140 mM NaCl, 2mM KCl, 4.5 mM MgCl2, 1.5mM CaCl2, and 5mM HEPES-NaOH with pH = 7.1) was applied to the window and then the cuticle and connective tissue were dissected to expose the SEZ. Mounted flies were placed on a Nikon A1R confocal microscope and then imaged using a 20X water-dipping objective lens. The pinhole was opened to allow a thicker optical section to be monitored. All recordings were taken at 4Hz with 256 resolution. Similar to PER, tastants were applied to the proboscis for 1-2s with a wick, which was operated using a micromanipulator (Narishige International USA, Inc.; Amityville, NY). For analysis, regions of interest were drawn manually around posterior IR56D projections. Baseline fluorescence was calculated as the average fluorescence of the first 5 frames, beginning 10 sec prior to tastant application. For each frame, the % change in fluorescence (%ΔF/F) was calculated as: (peak fluorescence - baseline fluorescence)/baseline fluorescence * 100. Average fluorescence traces were created by taking the average and standard error of %ΔF/F for each recording of a specific tastant.

### Aversive Taste Memory

Taste discrimination was assessed my measuring aversive taste memory, as described previously(Tauber et al., 2017). Female flies were starved for 48 h prior to each experiment. Flies were then anaesthetized on CO_2_ and the thorax of each fly was glued to a microscope slide using clear nail polish (#451D; Wet n Wild, Los Angeles, CA). Flies were acclimated to these conditions in a humidified box for 60 min. For each experiment, the microscope slide was mounted vertically under a dissecting microscope (#SM-1BSZ-144S; AmScope; Irvine, California). Flies were water satiated prior to each experiment and in between each test/training session. For tastant presentation, we used a 200 μL pipet tip attached to a 3ml syringe (#14-955-457; Fisher Scientific). For the pretest, 1% fatty acid was presented to the proboscis 3 times, with 1 min in between each presentation, and the number of full proboscis extensions was recorded. During training, a similar protocol was used except that each tastant presentation was immediately followed by 50mM quinine presentation which flies were allowed to drink it for up to 2 sec or until an extended proboscis was retracted. A total of 3 training sessions were performed. In between each session, the proboscis was washed with water and flies were allowed to drink to satiation. To assess taste discrimination, flies were tested either with that same tastant without quinine or with an untrained tastant. Another group of flies were tested as described above but quinine was never presented (naïve). At the end of each experiment, flies were given 1M sucrose to check for retained ability to extend proboscis and all non-responders were excluded.

### Statistical Analysis

All measurements are presented as bar graphs showing mean ± standard error. Measurements of PER and aversive taste memory were not normally distributed and so the non-parametric Kruskal-Wallis test was used to compare two or more genotypes. To compare two or more genotypes and two treatments, a restricted maximum likelihood (REML) estimation was used. For data that was normally distributed (calcium imaging data), a one-way or two-way analysis of variance (ANOVA) was used for comparisons between two or more genotypes and one treatment or two or more genotypes and multiple treatments, respectively. All post hoc analyses were performed using Sidak’s multiple comparisons test. Statistical analyses and data presentation were performed using InStat software (GraphPad Software 8.0; San Diego, CA).

## Acknowledgments

We would like to thank members of the Keene and Dahanukar labs for technical assistance and helpful discussion. This work was supported by NIH grants R01 NS085252 to ACK and R01DC017390 to ACK and AD, as well as support from FAU’s Jupiter Life Science Initiative.

## References

Ahn, J. E., Chen, Y., & Amrein, H. (2017). Molecular basis of fatty acid taste in Drosophila. ELife, 6, 1–21. https://doi.org/10.7554/eLife.30115.001

Akerboom, J., Chen, T.-W., Wardill, T. J., Tian, L., Marvin, J. S., Mutlu, S., … Looger, L. L. (2012). Optimization of a GCaMP Calcium Indicator for Neural Activity Imaging. The Journal of Neuroscience, 32(40), 13819–13840. https://doi.org/10.1523/JNEUROSCI.2601-12.2012

Amin, H., & Lin, A. C. (2019). Neuronal mechanisms underlying innate and learned olfactory processing in Drosophila. Current Opinion in Insect Science. https://doi.org/10.1016/j.cois.2019.06.003

Benton, R., Vannice, K. S., Gomez-Diaz, C., & Vosshall, L. B. (2009). Variant Ionotropic Glutamate Receptors as Chemosensory Receptors in Drosophila. Cell, 136(1), 149–162. https://doi.org/10.1016/j.cell.2008.12.001

Cameron, P., Hiroi, M., Ngai, J., & Scott, K. (2010). The molecular basis for water taste in Drosophila. Nature, 465(7294), 91–95. https://doi.org/10.1038/nature09011

Carleton, A., Accolla, R., & Simon, S. A. (2010). Coding in the mammalian gustatory system. Trends in Neurosciences. https://doi.org/10.1016/j.tins.2010.04.002

Chaudhari, N., & Roper, S. D. (2010). The cell biology of taste. Journal of Cell Biology, 190(3), 285–296. https://doi.org/10.1083/jcb.201003144

Chen, Y.-C. D., & Dahanukar, A. (2020). Recent advances in the genetic basis of taste detection in Drosophila. Cellular and Molecular Life Sciences, 77(6), 1087–1101. https://doi.org/10.1007/s00018-019-03320-0

Chu, B., Chui, V., Mann, K., & Gordon, M. D. (2014). Presynaptic gain control drives sweet and bitter taste integration in Drosophila. Current Biology, 24(17), 1978–1984. https://doi.org/10.1016/j.cub.2014.07.020

Clyne, P. J., Warr, C. G., & Carlson, J. R. (2000). Candidate taste receptors in Drosophila. Science (New York, N.Y.), 287(5459), 1830–1834. https://doi.org/10.1126/science.287.5459.1830

Cognigni, P., Felsenberg, J., & Waddell, S. (2018). Do the right thing: neural network mechanisms of memory formation, expression and update in Drosophila. Current Opinion in Neurobiology, 49, 51–58. https://doi.org/10.1016/j.conb.2017.12.002

Dahanukar, A., Lei, Y.-T., Kwon, J. Y., & Carlson, J. R. (2007). Two Gr Genes Underlie Sugar Reception in Drosophila. Neuron, 56(3), 503–516. https://doi.org/https://doi.org/10.1016/j.neuron.2007.10.024

DasGupta, S., & Waddell, S. (2008). Learned Odor Discrimination in Drosophila without Combinatorial Odor Maps in the Antennal Lobe. Current Biology, 18(21), 1668–1674. https://doi.org/10.1016/j.cub.2008.08.071

Devineni, A. V., Sun, B., Zhukovskaya, A., & Axel, R. (2019). Acetic acid activates distinct taste pathways in Drosophila to elicit opposing, state-dependent feeding responses. ELife, 8, e47677. https://doi.org/10.7554/eLife.47677.001

Flood, T. F., Iguchi, S., Gorczyca, M., White, B., Ito, K., & Yoshihara, M. (2013). A single pair of interneurons commands the Drosophila feeding motor program. Nature, 499(7456), 83–87. https://doi.org/10.1038/nature12208

Freeman, E. G., & Dahanukar, A. (2015). Molecular neurobiology of Drosophila taste. Current Opinion in Neurobiology, 34, 140–148. https://doi.org/10.1016/j.conb.2015.06.001

Guven-Ozkan, T., & Davis, R. L. (2014). Functional neuroanatomy of Drosophila olfactory memory formation. Learning & Memory (Cold Spring Harbor, N.Y.), 21(10), 519–526. https://doi.org/10.1101/lm.034363.114

Hallem, E. A., & Carlson, J. R. (2006). Coding of Odors by a Receptor Repertoire. Cell, 125(1), 143–160. https://doi.org/10.1016/j.cell.2006.01.050

Hallem, E. A., Dahanukar, A., & Carlson, J. R. (2006). Insect odor and taste receptors. Annual Review of Entomology, 51(10), 113–135. https://doi.org/10.1146/annurev.ento.51.051705.113646

Harris, D. T., Kallman, B. R., Mullaney, B. C., & Scott, K. (2015). Representations of Taste Modality in the Drosophila Brain. Neuron, 86(6), 1449–1460. https://doi.org/10.1016/j.neuron.2015.05.026

Inagaki, H. K., Panse, K. M., & Anderson, D. J. (2014). Independent, reciprocal neuromodulatory control of sweet and bitter taste sensitivity during starvation in Drosophila. Neuron, 84(4), 806–820. https://doi.org/10.1016/j.neuron.2014.09.032

Jiao, Y., Moon, S. J., Wang, X., Ren, Q., & Montell, C. (2008). Gr64f Is Required in Combination with Other Gustatory Receptors for Sugar Detection in Drosophila. Current Biology, 18(22), 1797–1801. https://doi.org/https://doi.org/10.1016/j.cub.2008.10.009

Kang, K., Pulver, S. R., Panzano, V. C., Chang, E. C., Griffith, L. C., Theobald, D. L., & Garrity, P. A. (2010). Analysis of Drosophila TRPA1 reveals an ancient origin for human chemical nociception. Nature, 464(7288), 597–600. https://doi.org/10.1038/nature08848

Keene, A. C., & Masek, P. (2012). Optogenetic induction of aversive taste memory. Neuroscience, 222. https://doi.org/10.1016/j.neuroscience.2012.07.028

Keller, A., Gerkin, R. C., Guan, Y., Dhurandhar, A., Turu, G., Szalai, B., … Zupan, B. (2017). Predicting human olfactory perception from chemical features of odor molecules. Science, 355(6327), 820–826. https://doi.org/10.1126/science.aal2014

Kirkhart, C., & Scott, K. (2015). Gustatory learning and processing in the Drosophila. Journal of Neuroscience, 15(35), 5950–5958.

Koh, T. W., He, Z., Gorur-Shandilya, S., Menuz, K., Larter, N. K., Stewart, S., & Carlson, J. R. (2014). The Drosophila IR20a Clade of Ionotropic Receptors Are Candidate Taste and Pheromone Receptors. Neuron, 83(4), 850–865. https://doi.org/10.1016/j.neuron.2014.07.012

Kubrak, O. I., Lushchak, O. V., Zandawala, M., & Nässel, D. R. (2016). Systemic corazonin signalling modulates stress responses and metabolism in *Drosophila*. Open Biology, 6(11), 160152. https://doi.org/10.1098/rsob.160152

LeDue, E. E., Mann, K., Koch, E., Chu, B., Dakin, R., & Gordon, M. D. (2016). Starvation-Induced Depotentiation of Bitter Taste in Drosophila. Current Biology, 26(21), 2854–2861. https://doi.org/10.1016/j.cub.2016.08.028

Levis, R., Hazelrigg, T., & Rubin, G. M. (1985). Effects of genomic position on the expression of transduced copies of the white gene of *Drosophila*. Science (New York, N.Y.), 229(4713), 558–561. https://doi.org/10.1126/science.2992080

Mann, K., Gordon, M., & Scott, K. (2013). A pair of interneurons influences the choice between feeding and locomotion in Drosophila. Neuron, 79(4), 754–765. https://doi.org/10.1016/j.neuron.2013.06.018

Marella, S., Fischler, W., Kong, P., Asgarian, S., Rueckert, E., & Scott, K. (2006). Imaging taste responses in the fly brain reveals a functional map of taste category and behavior. Neuron, 49(2), 285–295. https://doi.org/10.1016/j.neuron.2005.11.037

Marella, S., Mann, K., & Scott, K. (2012). Dopaminergic Modulation of Sucrose Acceptance Behavior in Drosophila. Neuron, 73(5), 941–950. https://doi.org/10.1016/j.neuron.2011.12.032

Masek, P, Worden, K., Aso, Y., Rubin, G., & Keene, A. (2015). A dopamine-modulated neural circuit regulating aversive taste memory in Drosophila. Curr. Biol., 25(11), 1535–1541.

Masek, Pavel, & Keene, A. C. (2013). Drosophila Fatty Acid Taste Signals through the PLC Pathway in Sugar-Sensing Neurons. PLoS Genetics, 9(9). https://doi.org/10.1371/journal.pgen.1003710

Masek, Pavel, & Scott, K. (2010). Limited taste discrimination in Drosophila. Proceedings of the National Academy of Sciences of the United States of America, 107(33), 14833–14838. https://doi.org/10.1073/pnas.1009318107

Masek, Pavel, Worden, K., Aso, Y., Rubin, G. M., & Keene, A. C. (2015). A dopamine-modulated neural circuit regulating aversive taste memory in drosophila. Current Biology, 25(11), 1535–1541. https://doi.org/10.1016/j.cub.2015.04.027

Mishra, D., Thorne, N., Miyamoto, C., Jagge, C., & Amrein, H. (2018). The taste of ribonucleosides: Novel macronutrients essential for larval growth are sensed by Drosophila gustatory receptor proteins. PLoS Biology, 16(8), 1–19. https://doi.org/10.1371/journal.pbio.2005570

Nara, K., Saraiva, L. R., Ye, X., & Buck, L. B. (2011). A large-scale analysis of odor coding in the olfactory epithelium. Journal of Neuroscience, 31(25), 9179–9191. https://doi.org/10.1523/JNEUROSCI.1282-11.2011

Parnas, M., Lin, A. C., Huetteroth, W., & Miesenb??ck, G. (2013). Odor Discrimination in Drosophila: From Neural Population Codes to Behavior. Neuron, 79(5), 932–944. https://doi.org/10.1016/j.neuron.2013.08.006

Pfeiffer, B. D., Ngo, T. T. B., Hibbard, K. L., Murphy, C., Jenett, A., Truman, J. W., & Rubin, G. M. (2010). Refinement of tools for targeted gene expression in Drosophila. Genetics, 186(2), 735–755. https://doi.org/10.1534/genetics.110.119917

Pool, A. H., Kvello, P., Mann, K., Cheung, S. K., Gordon, M. D., Wang, L., & Scott, K. (2014). Four GABAergic interneurons impose feeding restraint in Drosophila. Neuron, 83(1), 164–177. https://doi.org/10.1016/j.neuron.2014.05.006

Root, C. M., Masuyama, K., Green, D. S., Enell, L. E., Nässel, D. R., Lee, C. H., & Wang, J. W. (2008). A Presynaptic Gain Control Mechanism Fine-Tunes Olfactory Behavior. Neuron, 59(2), 311–321. https://doi.org/10.1016/j.neuron.2008.07.003

Rytz, R., Croset, V., & Benton, R. (2013). Ionotropic Receptors (IRs): Chemosensory ionotropic glutamate receptors in Drosophila and beyond. Insect Biochemistry and Molecular Biology. https://doi.org/10.1016/j.ibmb.2013.02.007

Sánchez-Alcañiz, J. A., Silbering, A. F., Croset, V., Zappia, G., Sivasubramaniam, A. K., Abuin, L., … Benton, R. (2018). An expression atlas of variant ionotropic glutamate receptors identifies a molecular basis of carbonation sensing. Nature Communications, 9(1), 4252. https://doi.org/10.1038/s41467-018-06453-1

Scott, K. (2018). Gustatory Processing in Drosophila melanogaster. Annual Review of Entomology, 63(1), 15–30. https://doi.org/10.1146/annurev-ento-020117-043331

Scott, K., Brady, R., Cravchik, A., Morozov, P., Rzhetsky, A., Zuker, C., & Axel, R. (2001). A Chemosensory Gene Family Encoding Candidate Gustatory and Olfactory Receptors in Drosophila. Cell, 104(5), 661–673. https://doi.org/10.1016/S0092-8674(01)00263-X

Slone, J., Daniels, J., & Amrein, H. (2007). Sugar Receptors in Drosophila. Current Biology, 17(20), 1809–1816. https://doi.org/https://doi.org/10.1016/j.cub.2007.09.027

Sweeney, S. T., Broadie, K., Keane, J., Niemann, H., & Kane, C. J. O. (1995). Targeted Expression of Tetanus Toxin Light Chain in Drosophila Specifically Eliminates Synaptic Transmission and Causes Behavioral Defects, 14, 341–351.

Tauber, J. M., Brown, E. B., Li, Y., Yurgel, M. E., Masek, P., & Keene, A. C. (2017). A subset of sweet-sensing neurons identified by IR56d are necessary and sufficient for fatty acid taste. PLOS Genetics, 13(11), e1007059. https://doi.org/10.1371/journal.pgen.1007059

Thakur, D., Kim, Y., Poudel, S., Montell, C., & Lee, Y. (2017). Calcium Taste Avoidance in Drosophila. Neuron, 97(1), 67–74.e4. https://doi.org/10.1016/j.neuron.2017.11.038

Thorne, N., Chromey, C., Bray, S., & Amrein, H. (2004). Taste Perception and Coding in Drosophila. Current Biology, 14(12), 1065–1079. https://doi.org/10.1016/j.cub.2004.05.019

van Giesen, L., & Garrity, P. A. (2017). More than meets the IR: the expanding roles of variant Ionotropic Glutamate Receptors in sensing odor, taste, temperature and moisture. F1000Research, 6(0), 1753. https://doi.org/10.12688/f1000research.12013.1

Vosshall, L. B., & Stocker, R. F. (2007). Molecular architecture of smell and taste in Drosophila. Annual Review of Neuroscience, 30, 505–533. https://doi.org/10.1146/annurev.neuro.30.051606.094306

Wang, Z., Singhvi, A., Kong, P., & Scott, K. (2004). Taste Representations in the Drosophila Brain. Cell, 117(7), 981–991. https://doi.org/10.1016/j.cell.2004.06.011

Wisotsky, Z., Medina, A., Freeman, E., & Dahanukar, A. (2011). Evolutionary differences in food preference rely on Gr64e, a receptor for glycerol. Nature Neuroscience, 14(12), 1534–1541. https://doi.org/10.1038/nn.2944

Yapici, N., Cohn, R., Schusterreiter, C., Ruta, V., & Vosshall, L. B. (2016). A Taste Circuit that Regulates Ingestion by Integrating Food and Hunger Signals. Cell, 165(3), 715–729. https://doi.org/10.1016/j.cell.2016.02.061

Yarmolinsky, D. A., Zuker, C. S., & Ryba, N. J. P. (2009). Common Sense about Taste: From Mammals to Insects. Cell. https://doi.org/10.1016/j.cell.2009.10.001

Zhang, Y. V., Ni, J., & Montell, C. (2013). The Molecular Basis for Attractive Salt-Taste Coding in Drosophila. Science, 340(6138), 1334–1338. https://doi.org/10.1126/science.1234133

Zhang, Yifeng, Hoon, M. A., Chandrashekar, J., Mueller, K. L., Cook, B., Wu, D., … Ryba, N. J. P. (2003). Coding of sweet, bitter, and umami tastes: Different receptor cells sharing similar signaling pathways. Cell, 112(3), 293–301. https://doi.org/10.1016/S0092-8674(03)00071-0

